# Emphasis on peripheral vision is accompanied by pupil dilation

**DOI:** 10.1101/2022.07.14.500035

**Authors:** Ana Vilotijević, Sebastiaan Mathôt

## Abstract

People are best able to detect stimuli in peripheral vision when their pupils are large, and best able to discriminate stimuli in central vision when their pupils are small. However, it is unclear whether our visual system makes use of this by dilating the pupils when attention is directed towards peripheral vision. Therefore, we tested whether pupil size adapts to the ‘breadth’ of attention. We found that pupils dilate with increasing attentional breadth, both when attention is diffusely spread and when attention is directed at specific locations in peripheral vision. We further found a correlation with performance, suggesting a functional benefit of this effect. Based on our results and others, we propose that cognitively driven pupil dilation is not an epiphenomenal marker of Locus Coeruleus activity, as is often assumed, but rather is an adaptive response that reflects an emphasis on peripheral vision.

## Introduction

Spatial attention is commonly seen as a spotlight that highlights a certain region in space and thus enhances processing for that region. Importantly, the size of this spotlight is not constant, but adapts to the demands of the situation (Eriksen & St. James, 1986; Greenwood & Parasuraman, 1999, 2004; Lawrence et al., 2020; Müller et al., 2003). For example, when trying to find a certain figure in a book, your attention is broadly distributed as you are flipping through the pages. Yet, once you have found the figure, your attention narrows and focuses on the details of the figure and the caption underneath. The zoom-lens theory of attention was developed to accommodate the fact that the focus of spatial attention not only has a location but also a size (Eriksen & St. James, 1986; Eriksen & Yeh, 1985; LaBerge, 1983). A change in the size of this zoom lens is commonly referred to as a change in *attentional breadth*, which gets bigger when attention is diffusely spread across the visual field, thus also encompassing peripheral vision, and smaller when attention is focused centrally, encompassing only (para)foveal vision (Brocher et al., 2018; Mathôt, 2020).

The zoom-lens theory postulates an inverse relationship between the breadth of attention and visual processing quality; in other words, broadly distributed attention results in parallel but superficial processing of stimuli, while narrowly distributed attention allows for detailed scrutiny of stimuli. Referring back to the example from the beginning, broadly distributed attention allows us to quickly scan the pages until we find the right figure, based on a general impression of what this figure should look like, but at the expense of fine resolution for the text presented on those pages. Conversely, narrowly distributed attention provides us with sharp and detailed information about the attended location, but at the expense of sensitivity for things appearing in the periphery.

These functional differences in processing quality between broadly and narrowly focused attention map onto physiological differences between foveal and peripheral vision at the level of the retina. Firstly, photoreceptors are not uniformly distributed across the retina: the density of cones is highest near the fovea, and declines toward the periphery (Hirsch & Curcio, 1989; Pumphrey, 1948; Rosenholtz, 2016). Furthermore, in the fovea, photoreceptors have one-to-one connections to ganglion cells, which allows photoreceptor input to be transmitted with minimal loss of spatial information. In contrast, in the peripheral retina, many photoreceptors (both rods and cones) project to a single shared ganglion cell, thus losing spatial information. Because of these physiological properties of the retina, foveal vision provides higher visual acuity than peripheral vision does (Wolfe et al., 2009; Curcio et al., 1987).

The asymmetry between peripheral and foveal vision also has implications for the optimal size of the pupil. When the pupil is constricted, only a small part of the lens is exposed, thus lessening optical distortions (both spherical and chromatic) that cause optical blur (Mathôt, 2018; Wang & Ciuffreda, 2006), in turn resulting in high visual acuity. In contrast, when the pupil is dilated, a larger part of the lens is exposed, thus allowing more light to enter the eye, in turn resulting in increased visual sensitivity (see also Franke et al., 2022 for related effects of pupil size on retinal sensitivity in mice). The size of the pupil therefore reflects a trade-off between visual acuity and sensitivity, where small pupils favor acuity over sensitivity. Crucially, due to the physiology of the retina, high visual acuity (and thus a small pupil) is mainly beneficial for foveal vision, whereas it is largely wasted on peripheral vision.

The notion that large pupils are beneficial for peripheral vision has received empirical support from a study by Mathôt & Ivanov (2019) who manipulated pupil size while measuring performance on a peripheral detection task and on a central discrimination task. They found that detection of faint stimuli in peripheral vision is better with large pupils, whereas discrimination of fine-grained stimuli in foveal vision is better with small pupils. However, in this study, pupil size was manipulated, thus leaving open the question of whether our pupils also automatically adapt their size to the situation. More specifically, does pupil size increase with increasing attentional breadth to optimize perception for peripheral vision (and especially peripheral detection)?

This question has been addressed already in a number of studies (Brocher et al., 2018; Daniels et al., 2012; Klatt et al., 2021; Kolnes et al., 2022); for example, in a series of experiments conducted by Brocher and colleagues (2018; Klatt et al., 2021), participants were exposed to a varying number of target stimuli presented laterally at various levels of eccentricity (12.5°, 27.5°, and 42.5°) relative to a central fixation dot. Prior to the presentation of the targets, participants received a cue that informed them of the eccentricity of the upcoming targets; this served as a manipulation of attentional breadth. Finally, participants reported the number of targets on each side of the display. The results showed that pupil size increased with increasing attentional breadth. However, task performance was poorer at the far-(42.5°) and medium-eccentricity (27.5°) conditions as compared to the near-eccentricity condition (12.5°); this makes it impossible to disentangle whether the increase in pupil size with increasing attentional breadth simply originated from greater mental effort (because the task was more difficult for the medium and far eccentricities), or from attentional breadth per se. Daniels and colleagues (2012) tested the same hypothesis in a set of experiments in which participants were presented with stimuli in a diamond-like configuration; stimuli were displayed centrally and/or peripherally. Participants covertly attended either to the central (narrow attention) or peripheral (broad attention) stimuli based on a cue. For example, in one experiment, the luminance of a central dot changed on a trial-by-trial basis, indicating narrow or broad attention. Even though their results suggested that pupil size increased with increasing attentional breadth, it was not mentioned if the cues were counterbalanced, which is important in order to disentangle the effect of visual stimulation on pupil responses from the effect of attentional breadth per se. Finally, in a very recent study, Kolnes and colleagues (2022) manipulated attentional breadth by cueing participants’ attention either to a small or a large circular rim at different eccentricities (near and far), and asked them to remember the location of a gap in the cued rim. Consistent with the above-mentioned studies, they found that pupil size increased with increasing attentional breadth (i.e., when the large rim was cued). Even though Kolnes and colleagues (2022) aimed to carefully control for common confounds (visual input and gaze position), it remained unclear if auditory cues, used to induce different levels of attentional breadth, were counterbalanced; again this is crucial because not only visual, but also auditory, stimuli have an effect on the pupil (Gingras et al., 2015). In addition, task difficulty was not controlled between conditions, although in this case the task was easier, rather than more difficult, for the far eccentricity as compared to the near eccentricity (i.e., opposite to the paradigm used by Brocher et al., 2018; Klatt et al. 2021).

Taken together, several studies have provided preliminary evidence for the effect of attentional breadth on pupil size, such that pupil size increases with increasing attentional breadth; however, none of these studies simultaneously controlled for all potential confounds. Therefore, in the current study, we tested the same hypothesis while carefully controlling for all potential confounds. We believe that this is crucial because of the theoretical importance of the relationship between attentional breadth and pupil size.

In addition, we explored different operationalizations of attentional breadth, which in previous studies have not been clearly dissociated. Specifically, we distinguish three theoretically possible forms of attention. First, the attentional spotlight can grow or shrink in the shape of a filled circle that is centered on the point of fixation, such that attention is never disengaged from central vision; in this view, increased attentional breadth is similar to a general increase in vigilance for anything that may appear at any location (Figure 1a). Second, the attentional spotlight can grow or shrink in the shape of an annulus (ring); in this view, increased attentional breadth is also similar to a general increase in vigilance, but with the added assumption that we either attend to (a specific eccentricity in) peripheral vision, or to central vision, but not to both (Figure 1b). Third, the attentional spotlight can be directed towards a single location in central or peripheral vision; this kind of attention is similar to spatial attention as manipulated in traditional cueing paradigms, and in this view increased attentional breadth is similar to a spatial shift of attention towards a location in the periphery (Figure 1c). All of these different types of attention allow for an increased or decreased emphasis on peripheral vision, but the specific way in which this happens differs.

**Figure 1.**
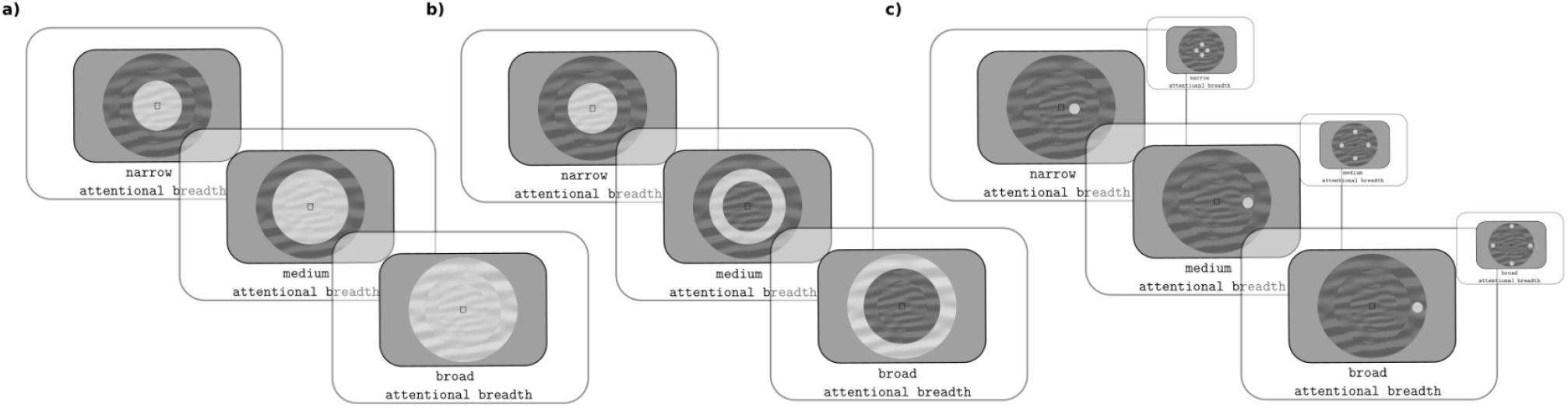
Different operationalizations of attentional breadth. *(a)* Attentional spotlight in the shape of a filled circle. *(b)* Attentional spotlight in the shape of an annulus. *(c)* Attentional spotlight directed towards a single location in central or peripheral vision.

Finally, we explored the relationship between attentional breadth, pupil size, and behavioral performance, which is important to better understand the role of pupil size in behavior.

## Results

To investigate the effect of attentional breadth on pupil size, we analyzed eye-tracking data from 100 participants performing a visual discrimination task. We conducted three separate experiments, which are described in detail under Materials and methods and analyzed separately as described in the Supplemental Information; however, because the task and design of all three experiments were very similar, and we wanted to test the interaction with the type of attentional breadth, we will focus on the combined data below. (The experiments were preregistered separately. The combined analysis was not preregistered.)

Participants saw three concentric annuli that occupied various eccentricities (near [1.16°], medium [3.47°], and far [10.40°]), corresponding to three sizes of attentional breadth. Next, after an interval of at least 2000 ms, a target appeared within one of the annuli; the target was either a slight increment or decrement of luminance relative to the background, and participants reported its identity (luminance increment or decrement). To avoid differences in task difficulty between eccentricities, we used an interleaved staircase procedure to keep accuracy constant across eccentricities by varying the visibility of the target. Crucially, 1000 ms before the presentation of the annuli, we cued participants’ covert attention by indicating either the annulus (attentional spotlight’s Size Condition) or the specific location within the annulus (attentional spotlight’s Location Condition) where the upcoming target would most likely (but not always) appear (see Figure 2a). The Attentional Breadth Type (Size Condition vs Location Condition) was partly varied between participants (Exp 1: Size; Exp 2: Location) and partly within participants (Exp 3: Size and Location, counterbalanced between blocks). Participants were required to keep their gaze fixated in the middle of the screen throughout the trial.

**Figure 2.**
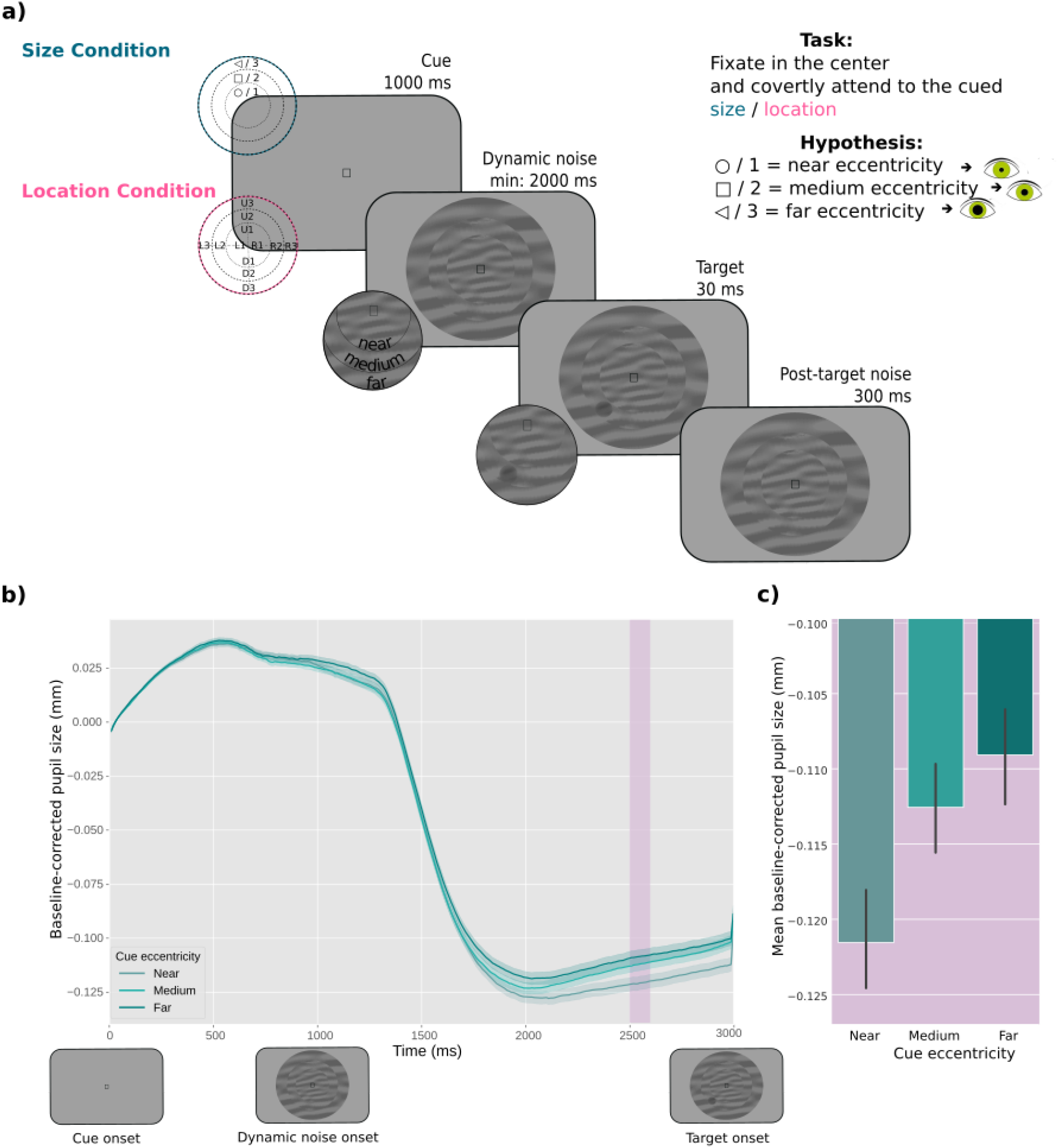
*a)* Schematic trial sequence. In the Size Condition, participants’ attention was cued to one of the differently sized annuli. In the Location Condition, participants’ attention was cued to a specific location within one of the annuli. Participants reported whether a target was a luminance increment or, as in this example, a luminance decrement embedded in a dynamic stream of noise. *b)* The effect of attentional breadth on pupil size. The y-axis represents baseline-corrected pupil size in millimeters. The x-axis represents time in milliseconds since cue onset. The panels below the x-axis represent the order of the events in the experiments.

Differently colored lines represent the three cue eccentricities (near, medium, far) corresponding to different levels of attentional breadth. The pink-shaded area represents the time window (2500-2600ms) where the effect was the strongest, as determined through a cross-validation procedure. Error bands indicate the grand standard error. *c)* Mean baseline-corrected pupil size in millimeters as a function of cue eccentricity within the selected time window (2500-2600ms). Error bars indicate the grand standard error.

### The effect of attentional breadth on pupil size

We ran a four-fold cross-validation analysis on a predetermined time-window of 750-3000 ms after the onset of the cue to assess differences in pupil size across eccentricities. This procedure is described in more detail in Mathôt and Vilotijević (2022); however, in brief, this analysis determines a time point at which an effect of interest (Cue Eccentricity in our case) is strongest by splitting the data into training and test sets. Next, for each test set, the corresponding training set is used to determine the time point at which the effect is the strongest; a single linear mixed effects (LME) analysis is then conducted for the full dataset, using for each test set the time point that was selected based on the corresponding training set, thus avoiding both circularity and the need to correct for multiple comparisons. This means that the dependent variable consists of a column of (baseline-corrected) pupil-size values that can (and often do) correspond to different samples for different trials.

Our model included pupil size as dependent variable, Cue Eccentricity as fixed effect (coded ordinally: −1 = *near*, 0 = *medium*, 1 = *far*; this was the case for subsequent analyses as well), and by-participant random intercepts and slopes. The lower boundary of our period of interest (750 ms) was decided on the basis of the finding that a voluntary shift of covert visual attention towards a bright or dark surface affected pupil size from about 750 ms after cue onset (Mathôt et al., 2013). The upper boundary (3000 ms) was set to the first moment at which the target could appear in any of the three experiments. The effect of Cue Eccentricity on pupil size emerged around 1750 ms after cue onset and the cross-validation analysis showed that it reached maximum around 2500 and 2600 ms after cue onset (*t* = 2.75, *p* = .006). Specifically, and in line with our hypothesis, we found that the pupil size increased with increasing attentional breadth (see Figure 2b, c).

Next, to test whether the effect of attentional breadth differed between the Size Condition and Location Condition, we focused on mean pupil size during the 2500-2600 ms window as selected by the cross-validation analysis. We conducted an LME analysis with mean pupil size during the selected interval as dependent variable, Cue Eccentricity and Attentional Breadth Type (reference: Size Condition) as fixed effects, and by-participant random intercepts (a model with by-participant random slopes failed to converge). This revealed, as before, a main effect of Cue Eccentricity (*b* = 0.008, SE = 0.003, *t* = 2.77, *p* = .006), but no main effect of Attentional Breadth Type (*b* = 0.008, SE = 0.005, *t* = 1.58, *p* = .115), and—more importantly—no interaction between Cue Eccentricity and Attentional Breadth Type (*b* = −0.003, SE = 0.004, *t* = 0.74, *p* = .457).

Taken together, we found an effect of attentional breadth on pupil size, but this effect did not reliably depend on the type of attentional breadth (Size Condition vs Location Condition).

### Behavioral cueing effect

Next, to test the behavioral cueing effect, we ran Linear Mixed Models (LMM) testing a model that included Accuracy as dependent variable, Cue Validity as fixed effect, and by-participant random intercepts and slopes for Cue Validity. The results showed a behavioral cueing effect; that is, participants were more accurate on valid versus invalid trials (*b* = 0.17, SE = 0.03, *z* = 5.02, *p* < .001; Figure 3a).

**Figure 3.**
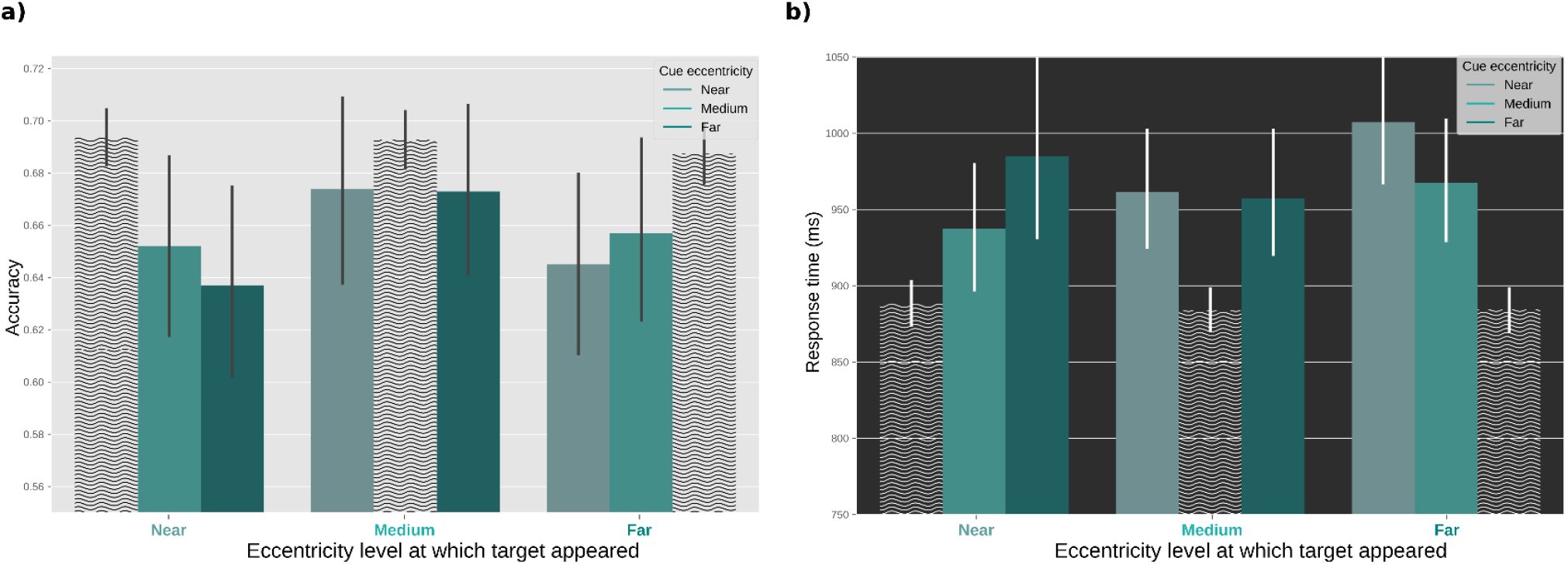
Behavioral results. a) Accuracy as a function of the eccentricity at which the target appeared (*x*-axis) and the eccentricity that was cued (bars). b) Reaction times as a function of the eccentricity at which the target appeared (*x*-axis) and the eccentricity that was cued (bars). Shaded wavy bars represent valid conditions. Error bars represent the grand standard error.

### Task difficulty

As explained in the Methods section, the accuracy across conditions was controlled by a staircase procedure, keeping it at around 70% for validly cued trials, to rule out the potential effect of mental effort. However, participants may have still perceived some conditions to be more difficult than others, and consequently responded more slowly to those trials. To check for this, we ran a Linear Mixed Model (LMM) testing a model that included Reaction Times as dependent variable, Cue Eccentricity as fixed effect, and by-participant random intercepts and slopes for Cue Eccentricity. The results showed that there were no significant differences in RT across conditions (*b* = −2.68, SE = 4.20, *z* = −.64, *p* = .523; Figure 3b). In other words, there was no difference in task difficulty between different levels of attentional breadth, neither in terms of accuracy nor reaction times.

### Correlations between pupil size and behavior

Figure 4a depicts individual differences across participants in the strength of the effect of attentional breadth on pupil size; here, the “Attentional Breadth” effect corresponds to the Pearson correlation coefficient between Cue Eccentricity and mean pupil size during the 2500-2600 ms window, determined for each participant separately. About two-thirds of the participants showed an effect in the hypothesized direction; in other words, although the effect of attentional breadth is highly reliable at the group level, there is a lot of interindividual variability.

**Figure 4.**
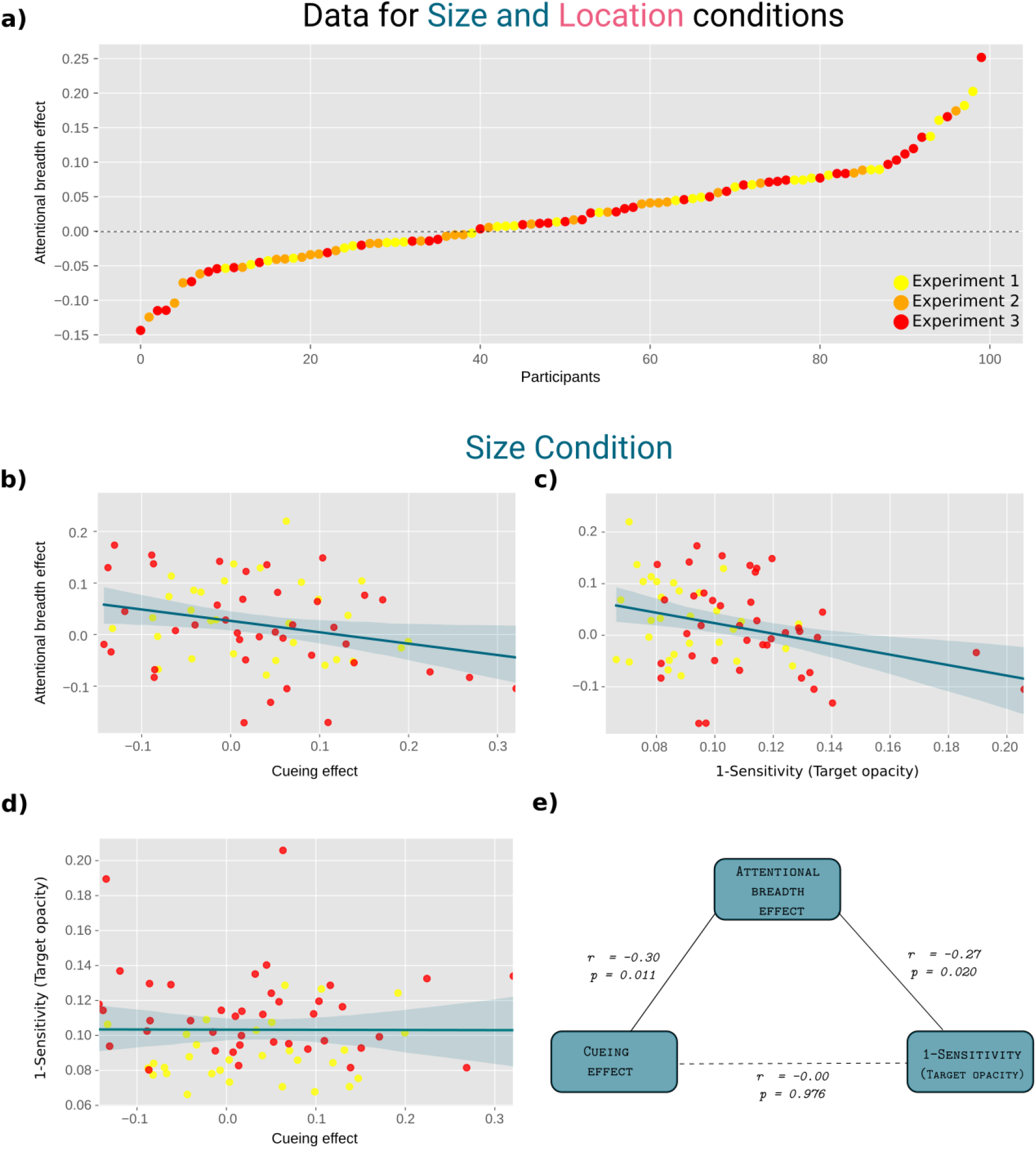
*a)* Attentional-breadth effect sizes for individual participants, sorted by effect size for both the Size and Location Conditions. *b)* Correlation between attentional-breadth effect size and behavioral cueing effect (Size Condition only). *c)* Correlation between attentional-breadth effect size and inverse sensitivity measure (target opacity; Size Condition only). *d)* Correlation between inverse sensitivity and cueing effect (Size Condition only). *e)* Schematic illustration of the link between attentional breadth effect, cueing effect, and inverse sensitivity with corresponding correlations (Size Condition only). *Note*: Detection of targets with lower opacities meant higher sensitivity.

**Figure 5.**
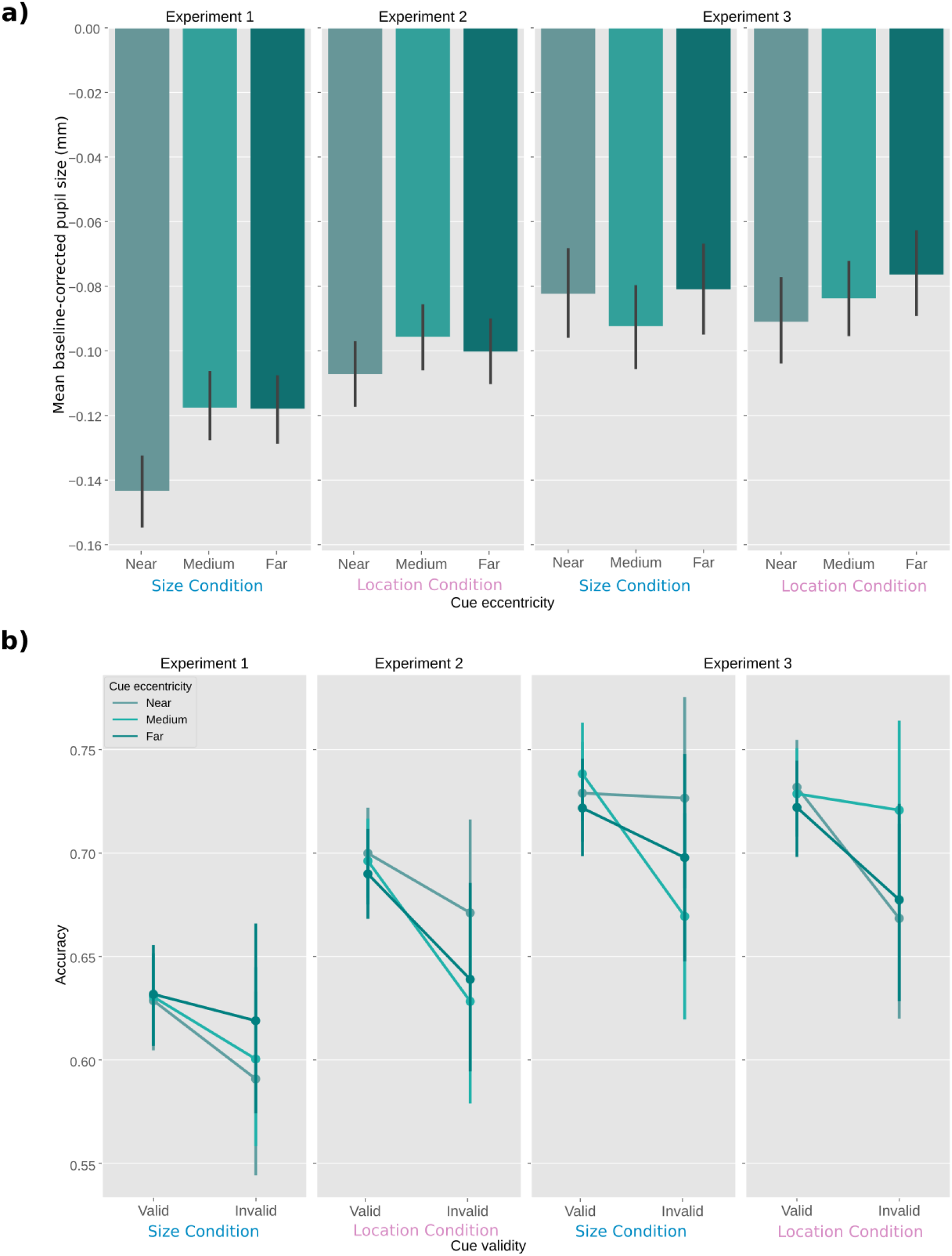
*a)* Mean baseline-corrected pupil size expressed in millimeters as a function of cue eccentricity across experiments. *b)*. Accuracy as a function of cue validity across experiments and corresponding conditions. Error bars indicate 95% confidence interval.

Next, we assessed how the attentional-breadth effect relates to visual sensitivity (Figure 4c); here, sensitivity corresponds to 1 - Target Opacity, because the staircase procedure converged on a low target opacity for participants who performed well. We found a significant negative correlation between the attentional-breath effect size and Target Opacity. In other words, participants who exhibited a stronger effect of attentional breadth on pupil size were better able to detect targets whose opacity was low (higher task difficulty). However, this was only true for the Size Condition (*r* = −0.27, *p* = .020), and not for the Location Condition (*r* = .01, *p* = .957). These two correlations differed significantly (*z* = −1.99, *p* = .023).

We also assessed how the attentional-breadth effect relates to the behavioral cueing effect (Figure 4b); here, the cueing effect is calculated as the accuracy on valid trials minus the accuracy on invalid trials, determined for each participant separately. Strikingly—and unexpectedly—the attentional-breadth effect was negatively correlated with the behavioral cueing effect. In other words, participants who exhibited a stronger attentional-breadth effect exhibited a *weaker* behavioral cueing effect. However, this was again only true for the Size Condition (*r* = −.30, *p* = .011), and not for the Location Condition (*r* = .11, *p* = .380). (We will return to this in the Discussion.)

Finally, performance does not relate to the behavioral cueing effect (*r* = −.00, *p* = .976); that is, there is no systematic relationship between how well participants perform the task, and how strongly they show a behavioral cueing effect (Figure 4d).

## Discussion

We report three experiments that investigated whether pupil size increases with increasing attentional breadth, and if this effect depends on the type of attentional breadth, that is, whether attentional breadth is induced through changes in the *size* of the attentional spotlight (see Figure 1a,b), or changes in the *location* of the attentional spotlight (see Figure 1c). Large pupils are known to benefit detection of faint stimuli in peripheral vision (Mathôt, 2020; Wang et al., 2021; Woodhouse, 1975); therefore, the question of whether pupils reflexively dilate in response to increased attentional breadth is crucial for a better understanding of pupil responses as a form of ‘sensory tuning’: a subtle adjustment of the eyes to optimize visual information intake for a given situation (Mathôt, 2020; see also Franke et al., 2022).

To test this, participants had to discriminate the luminance of a target that was presented within one of three annuli that occupied various eccentricities (Figure 1a). We cued participants’ covert attention to either the annulus (Size Condition) or to the specific location within the annulus (Location Condition) where the upcoming target would most likely (but not always) appear. We measured pupil size in anticipation of the target. In doing so, we made sure that task difficulty (reflected both in accuracy and RTs), gaze position, and visual input (including luminance and contrast)—factors that are known to influence pupil size—were kept constant across conditions, which is a crucial improvement over previous studies that addressed this question (Brocher et al., 2018; Daniels et al., 2012; Klatt et al., 2021; Kolnes et al., 2022).

We found that pupil size increases with increasing attentional breadth. As shown in Figure 2b, the pupil slightly dilates during the cue presentation (1000 ms) after which it rapidly constricts once dynamic noisy annuli are shown; this constriction reflects the typical pupillary response to heavy visual stimulation (such as dynamic noise) and thus is consistent across conditions. Next and most importantly, pupil traces begin to differ between conditions around 1750 ms after the cue onset, reaching a maximum difference around 2500-2600 ms, and lingering until the target’s presentation. This means that when the target was expected to appear at the near level of attentional breadth (encompassing central vision), pupils were smaller than when the target was expected at the far or medium levels of attentional breadth (encompassing peripheral vision). Additionally, our data demonstrate that this effect does not, or hardly, depend on the type of attentional breadth, that is, on whether participants attend to entire circles at different eccentricities (Size Condition) or to specific locations at different eccentricities (Location Condition).

We further found that, between participants, sensitivity was reliably correlated with the effect of attentional breadth on pupil size, at least in the Size Condition (Figure 4c); that is, those participants whose pupils dilated strongly when attention was directed to the periphery were the same participants who performed well on the task. This supports the notion that tuning pupils to best serve peripheral and central vision enhances overall sensitivity, suggesting a functional benefit of attentional breadth.

Surprisingly, however, the behavioral cueing effect was negatively correlated with the effect of attentional breadth on pupil size (Figure 4b); that is, those participants whose pupil size was strongly affected by the cue were the same participants whose behavior was weakly affected by the cue. Possibly, this seemingly paradoxical pattern of results is an artifact of the task that we used in the current study. Specifically, participants needed to distinguish a subtle luminance increment from a decrement, which is a task that relies more on visual sensitivity than on acuity *also* when stimuli are presented in (para)foveal vision. To understand how this can result in a negative correlation between the behavioral cuing effect and the effect of attentional breadth on pupil size, imagine a hypothetical participant who shows a strong attentional breadth effect, that is, a participant whose pupils constrict strongly when the near annulus is cued. Across the board, this participant will perform relatively poorly when the near annulus is cued (because of the small pupils), even when the cue is valid and the target actually appears in the near annulus. Conversely, this participant will perform relatively well when the far or medium annulus is cued (because of larger pupils), even when the cue is invalid and the target actually appears in the near annulus. Because of this, the attentional breadth effect may, in our task, counteract the behavioral cuing effect. In turn, this may have resulted in a negative correlation between both effects. Importantly, in real-life—and in contrast to our experiment—tasks that people perform with (para)foveal vision, such as reading, generally emphasize visual acuity over sensitivity, whereas tasks that people perform with peripheral vision, such as detecting unexpected movement, generally emphasize visual sensitivity over acuity. An important avenue for future research will be to use laboratory tasks that more closely mimic how people use (para)foveal and peripheral vision in real life, and to investigate the functional role of the relationship between attentional breadth and pupil size in such tasks.

Although this was not the primary focus of our study, our behavioral results also speak to the question of whether, when an entire circle is cued (our Size Condition), attention takes the form of a filled circle (Fig. 1a) or of a ‘doughnut’/annulus (Fig. 1b). That is, are participants able to attend exclusively to the cued circle without also attending to the smaller circles within the cued circle (e.g., attending to the outermost circle in our stimulus display without also attending to the medium and innermost circles)? The answer appears to be yes: we found that attention was *selective* for the cued circle (Nobre & Kastner, 2014), as participants were consistently better performing on validly cued trials across *all* eccentricities (Figure 2). This suggests that attention was deployed in the shape of differently sized annuli, as depicted in Fig. 1b, rather than in the shape of differently sized spotlights/filled circles, as depicted in Fig. 1a. This is noteworthy because it implies an important difference between the concepts of attentional breadth and zoom-lens theory; the latter assumes attentional deployment in a form of differently sized filled circles (this also matches the concept of “attentional scaling” proposed by Lawrence et al., 2020), whereas attentional breadth assumes that attention is deployed in the form of differently sized annuli (Jefferies & Di Lollo, 2015; this also matches the concept of “doughnut” attention; Muller & Hubner, 2002). Our results suggest that attention can be distributed in the shape of a doughnut.

Finally but importantly, our results have implications for the Adaptive Gain Theory (AGT; Aston-Jones & Cohen, 2005). This theory differentiates between two modes of behavior—*exploitation* and *exploration*—and relates them to locus coeruleus (LC) activity and task utility (i.e. how potentially rewarding it is to perform a certain task). Specifically, information about task utility would be sent from higher-order brain structures (anterior cingulate cortex, orbitofrontal cortex) to LC, driving two modes of LC activity: phasic and tonic (Aston-Jones & Cohen, 2005; Usher et al., 1999). Phasic activity would reflect on-task behavior (exploitation), while tonic activity would reflect disengagement from the current task and seeking other tasks (exploration). Crucially, pupil size correlates with LC activity, and is therefore often used as a non-invasive measure of the extent to which people are in an exploration mode of behavior (large pupils) versus an exploitation mode of behavior (medium-to-small pupils) (Brink et al., 2016; Gilzenrat et al., 2010; Jepma & Nieuwenhuis, 2011). Our results add to this theory by offering a functional explanation for *why* there is a correlation between pupil size and modes of behavior. Specifically, an exploration mode of behavior is likely characterized by an emphasis on peripheral vision, and through this route may lead to pupil dilation. The same logic applies to other situations that are associated with pupil dilation. For example, high arousal is associated with pupil dilation (and thus high LC tonic activity mode; Binda & Murray, 2015; Bradley et al., 2008); thus far, arousal-related pupil dilation has been interpreted as an epiphenomenal physiological response. However, we propose that arousal-related pupil dilation is a way in which the visual system tunes itself to best serve the demands of the situation; specifically, in a situation that elicits high arousal (e.g., when you are afraid) it is crucial to be able to detect things anywhere in the visual field with little concern for their details; thus, the visual system recognizes that detection is more important than discrimination, and pupils consequently dilate to provide greater sensitivity.

The notion that small changes in pupil size can have substantial effects on visual processing at the level of the retina—a controversial notion among psychologists and vision scientists—has recently found strong support from a study involving mice (Franke et al., 2022); however, the mouse visual system differs in many ways from that of humans, and it is therefore important to work towards a better understanding of how pupil size interacts with the distribution and spectral sensitivity of photoreceptors to shape the early stages of visual processing in humans.

In sum, we demonstrate that pupil size increases with covert shifts of attention to the peripheral visual field, that is, with increasing attentional breadth, with no clear difference between different ways of deploying attention. We further found that participants who showed a large attentional-breadth effect on pupil size also performed better on the task, at least for the Size Condition, suggesting a functional benefit of the link between attentional breadth and pupil size. Finally, we propose that, more generally, cognitively driven pupil dilation reflects an emphasis on peripheral vision over foveal vision, and that this may explain why the pupil dilates in response to increased arousal or a switch to an exploration mode of behavior as postulated by the AGT.

## Limitations of the study

All experiments were conducted in a typical laboratory setting, where the experiment was run on regular-size computer monitors. Therefore, attention was never spread across the full visual field, which is about 180° in most people and thus extends far beyond the boundaries of a regular monitor.

## Materials and methods

### General

We conducted three experiments in order to test whether pupil size increases with increased attentional breadth. Each experiment’s methods, hypotheses, data sampling and analysis plans were preregistered on the Open Science Framework. All data, analysis scripts, and supplementary materials are available at https://osf.io/4nrgb/. The experiments were approved by the Ethics review board of the Department of Psychology at the University of Groningen (study approval code: PSY-2122-S-0139).

### Participants

In total, 111 (*N*_exp1_ = 30; *N*_exp2_ = 32; *N*_exp3_ = 49) first-year psychology students from the University of Groningen gave informed consent to participate in the study for course credits. For each experiment, we determined a target sample size and data exclusion criteria beforehand; we aimed for a target sample size of 30 participants for Experiment 1, and the same sample size was estimated for Experiment 2 based on a bootstrap power analysis conducted on the data from the first experiment (power = .81). Since we wanted to investigate an interaction effect in Experiment 3, and since we did not have a reliable effect-size estimate for a power analysis, we aimed for a target sample size of 40 participants. Prerequisites for retention of participant’s data were: 1) full completion of the experiment, 2) efficacy of the staircase procedure (see Data Exclusion). All participants had normal or corrected-to-normal vision. All participants were unique for each experiment.

### Apparatus and data acquisition

The experiments were programmed in OpenSesame (Mathôt et al., 2012) using PyGaze for eye tracking (Dalmaijer et al., 2014). Stimuli were presented on a 27-inch monitor (1920 × 1080 pixels resolution) and an Eyelink 1000 (sampling frequency of 1000 Hz; SR Research Ltd., Mississauga, Ontario, Canada), was used for eye tracking. Participants’ right eyes were recorded. All experiments were conducted in a dimly lit room.

### Procedure

Prior to the start of the experiment, the participant was well seated at about 60 cm distance from the computer monitor, with his or her chin placed on a chin-rest to keep the head in a stable position. First, a calibration-validation procedure was run. Before each trial, 1-point eye-tracker recalibration (“drift-correction”) was performed.

Throughout the experiments, participants were instructed to maintain their gaze in the center of the display, marked by a fixation dot. Each trial started with a centrally presented cue for 1000 ms that indicated where a target was most likely to appear. The cue display was followed by the dynamic noise display that consisted of concentric annuli that differed in their respective radii, occupying different eccentricities: near (1.16°), medium (3.47°), and far (10.40°). The annuli were filled with dynamic, oriented noise patches that changed every 30 Hz. The spatial frequency of the noise patches decreased with eccentricity, following the formula for cortical magnification as determined by Carrasco & Frieder (1997) for the upper visual field. At a random moment within the last second of the dynamic noise presentation, the target (a subtle luminance increment or decrement) was briefly flashed for 30 ms, after which the dynamic noise continued for another 300 ms. The exact time window in which the target appeared was 2000-3000 ms in Experiment 1 and 2500-3500 ms in Experiments 2 and 3. On a majority of trials (80%) the target appeared at the cued annulus or location (valid trials) while on the remaining 20% it appeared at a non-cued annulus or location (invalid trials). Participants’ task was to report whether the target was a luminance increment or decrement by pressing the left or right arrow on the keyboard, respectively. The sole difference between Experiment 1 and 2 was the type of the cue that was used, indicating the type of attentional breadth (Size or Location Condition). Specifically, in Experiment 1 we cued only the annulus (near, medium, or far) where the target was most likely to appear. To do so, we presented participants with symbolic cues (□, ○, ◁) that were indicative of each annulus. The mapping between cue symbol and eccentricity was counterbalanced across participants to ensure completely identical visual input in all conditions across participants. However, in Experiment 2, we cued both the annulus (near, medium, or far) and the location within the cued annulus (upper, lower, left, or right). Here, the cues were presented as a combination of codes for annuli (1/2/3 for near/medium/far) and codes for locations (U/D/R/L for up/down/right/left): for example, if U2 was presented that meant that the target was most likely to appear at the upper location of the middle annulus. This mapping was constant across all participants, because otherwise the task would be too complicated. Experiment 3 combined both cueing approaches (hence, both attentional breadth forms) in a blocked design, keeping the same cues as in Experiment 2 for the Location Condition, where a specific location within a specific annulus was cued (Location Condition), and using only numeric cues (1,2,3) for the condition where only an annulus was cued (Size Condition). All cues were the same size (.69°) and colored in black. All stimuli were superimposed on a gray background.

To ensure that the accuracy on validly cued trials was maintained at approximately 70%, we implemented a 2-up-1-down (Exp. 1) or 3-up-1-down (Exp. 2 and 3) staircase procedure (Leek, 2001), separately for each eccentricity, varying the opacity of the target with 1% steps. In case three correct responses were given in a row, the target opacity would be decreased by 1%, thus increasing task difficulty, while in case of a single incorrect response, the target opacity would be increased by 1%, thus decreasing task difficulty. Throughout the experiment, participants received feedback in the form of a black checkmark (.92°) or x mark (.92°), indicating correct and incorrect responses, respectively.

All experiments consisted of 330 trials that were shuffled and divided into 10 blocks. Experiment 1 and 2 consisted of 90 practice trials and 240 experimental trials. In Experiment 1, during practice trials, participants were shown both the cue in symbolic (□, ○, ◁) and written (‘near’,’medium’,’far’) form to make sure that they learned the mapping; between practice blocks, participants were asked to verbally repeat to the experimenter which symbol corresponds to which annulus, in order to verify that participants remembered the cue-annulus mapping. Experiment 3 consisted of 30 practice trials and 300 experimental trials, half of which represented Location Condition (5 successive blocks) and the other half represented Size Condition. The order of blocks was counterbalanced across participants.

### Data exclusion

Before analyzing the data, we checked if the staircase procedure was effective in keeping task difficulty equal across eccentricities. To that aim, after collecting the target sample size for each experiment, we compared participants’ performance when different eccentricities were cued by conducting Bayesian repeated measures ANOVA (with default parameters of JASP (JASP Team, 2022)) only on validly cued trials with Accuracy as dependent variable and Cue Eccentricity as independent variable. We used a Bayes Factor (BF_01_ > 3 as a threshold for finding substantial evidence in favor of the null hypothesis. This condition was not met in Experiment 3 (initial BF_01_ = 2.25), so we determined for each participant separately absolute performance deviance (this step was also preregistered), i.e., how much performance differed across eccentricities based on the following formula: `|Accuracy(near)-Accuracy(overall)| + |Accuracy(medium)-Accuracy(overall)| +|Accuracy(near)-Accuracy(overall)|`. Next, we iteratively excluded 9 participants with the largest absolute performance deviance until we found substantial evidence for the null hypothesis. Finally, we recruited additional participants to reach again the target sample size, after which we ended up with BF_01_ = 3.63. In Experiment 2, the staircase procedure did not work for one participant (*M*_accuracy_ = .44) while one other participant made a large number of eye movements (*GazeError_max_* = 19.07°). After these participants were replaced, we ended up with BF_01_ = 3.15. This condition was immediately met in Experiment 1 (BF_01_ = 6.29). Overall, the data from 100 participants was further analyzed (*N*_exp1_ = 30; *N*_exp2_ = 30; *N*_exp3_ = 40).

### Pupillary data: preprocessing

Following the workflow for preprocessing pupillary data that we described elsewhere (Mathôt & Vilotijević, 2022), we first interpolated blinks and downsampled the data by a factor of 10. Also, we converted pupil size measurements from arbitrary units to millimeters of diameter by using the formula specific to our lab (Wilschut & Mathôt, 2022). Next, we baseline-corrected the data by subtracting the mean pupil size during the first 50 ms after the onset of the cue (baseline period) from all subsequent pupil-size measurements on a trial-by-trial basis. Trials containing baseline pupil sizes of ± 2 *z*-scores were considered outliers, and hence excluded from the data. Missing trials (due to technical issues with recording in Experiment 2 and 3) were excluded as well (0.05%). In total, 1576 trials (Total = 5.91%; Exp1 = 5.58%, Exp2 = 5.44%, Exp3 = 6.51%) were excluded from the data.

## Additional information

## Acknowledgments

We thank Miran Pahlevan for his assistance in data collection and Hermine Berberyan for her valuable comments on the final manuscript.

## Author contribution

Conceptualization, Methodology, Software, Formal Analysis : A.V., S.M.; Investigation, Writing – Original Draft, Visualization: A.V.; Writing – Review & Editing: S.M., A.V.; Supervision, Funding Acquisition: S.M.

## Declaration of interests

The authors declare that there were no conflicts of interest with respect to the authorship or the publication of this article.

## Supplemental information

To test the attentional breadth effect in each experiment separately, we run LME with Mean Pupil Size during the selected time window (2500-2600 ms), determined by cross-validation analysis run on the collapsed data, as a dependent measure while Cue Eccentricity (ordinal: −1 = *near*, 0 = *medium*, 1 = *far*) was a fixed effect and by-participant random intercepts and slopes. Additionally, we tested cueing effect by conducting LMM using Accuracy as the dependent variable while Cue Validity was included as a fixed effect. The random-effect structure included a by-subjects random intercept and random slopes for Cue Validity.

*Experiment 1 - Size Condition*. The main effect of Cue Eccentricity was significant, *b* = 0.012, *SE* = 0.005, *t* = 2.696, *p* = 0.007. Looking at the behavioral performance, we observed a cueing effect, suggesting that participants were more accurate on valid trials, (*b* = 0.14, *SE* = 0.05, *z* = 2.61, *p* = 0.009).

*Experiment 2 - Location Condition*. When a specific location within a certain annulus was cued the pupillary effect did not reach significance (*b* = 0.004, *SE* = 0.004, *t* = 1.05, *p* = 0.315), although the pattern of results was still in line with overall results. The cueing effect was now even more prominent, *b* = 0.25, *SE* = 0.05, *z* = 4.63, *p* = 0.000.

*Experiment 3 - Size and Location Conditions*. The pupillary results were again in the hypothesized order, however, the results did not reach overall significance (*b* = 0.004, *SE* = 0.004, *t* = 1.21, *p* = 0.262). When analyzing conditions─Size Condition and Location Condition─separately, the pupillary results were in the hypothesized order, however, none of them reached significance: Size Condition: *b* = 0.001, *SE* = 0.005, *t* = 0.23, *p* = 0.821; Location Condition: *b* = 0.007, *SE* = 0.005, *t* = 1.45, *p* = 0.147. As regards the cueing effect, we observed the overall cueing effect in Experiment 3, *b* = 0.20, *SE* = 0.05, *z* = 4.31, *p* = 0.000). This was also true when only Location Condition was tested (*b* = 0.23, *SE* = 0.07, *z* = 3.42, *p* = 0.001) as well as for Size Condition (*b* = 0.18, *SE* = 0.07, *z* = 2.67, *p* = 0.008).

## Notes

https://osf.io/4nrgb/

### Competing Interest Statement

The authors have declared no competing interest.

### Summary of Updates

Figure 1 is added to clarify different operationalizations of attentional breadth.

